# Cardiac ventricular myosin and slow skeletal myosin exhibit dissimilar chemo-mechanical properties despite the same myosin heavy chain isoform

**DOI:** 10.1101/2022.03.10.483824

**Authors:** Tianbang Wang, Emrulla Spahiu, Florentine Behrens, Jennifer Osten, Fabius Grünhagen, Tim Scholz, Theresia Kraft, Arnab Nayak, Mamta Amrute-Nayak

**Affiliations:** Institute of Molecular and Cell Physiology, Hannover Medical School, 30625 Hannover, Germany

**Keywords:** Optical trapping, Single-molecule studies, actomyosin, cardiac ventricular myosin, skeletal myosin

## Abstract

The myosin II motors are ATP-powered, force-generating machines driving cardiac and muscle contraction. Myosin II heavy chain isoform-beta (β-MyHC) is primarily expressed in the ventricular myocardium and slow-twitch muscle fibers, such as in M. soleus. M. soleus-derived myosin II (SolM-II) is often used as an alternative to the ventricular β-cardiac myosin (βM-II); however, the direct assessment of detailed biochemical and mechanical features of the native myosins is limited. By employing the optical trapping method, we examined the mechanochemical properties of the native myosins isolated from rabbit heart ventricle and M. soleus muscles at the single-molecule level. Contrary to previous reports, the purified motors from the two tissue sources, despite the same MyHC isoform, displayed distinct motile and ATPase kinetic properties. βM-II was ∼threefold faster in the actin filament-gliding assay than SolM-II. The maximum acto-myosin (AM) detachment rate derived in single-molecule assays was ∼threefold higher in βM-II. The stroke size for both myosins was comparable. The stiffness of the ‘AM rigor’ cross-bridge was also similar for both the motor forms. The stiffness of βM-II was found to be determined by the nucleotide state of the actin-bound myosin. Our analysis revealed distinct kinetic differences, i.e., a higher AM detachment rate for the βM-II, corresponding to the ADP release rates from the cross-bridge, thus elucidating the observed differences in the motility driven by βM-II and SolM-II. These studies have important implications for the future choice of tissue sources to gain insights into cardiomyopathies

## Introduction

Cardiac contraction is driven by molecular motor proteins, myosin II. Two main cardiac-specific myosin II isoforms are expressed in higher mammalian heart namely, alpha myosin (αM-II) and beta myosin (βM-II). While in the atrium, primarily α-myosin is expressed, β-myosin is expressed in the left ventricle and interventricular septum. In the atrium, a mixture of both myosin isoforms is found, albeit with lower amounts of β-myosin. The distinct myosin isoform expression is linked to their activities, regulating the shortening speed of the different chambers, i.e., atrium and ventricle, as per their physiological requirement. Hundreds of cardiomyopathy causing mutations have been located in the ventricular myosin isoform (βM-II) responsible for varying degrees of severities in patients. Understanding the key functional characteristics of this myosin isoform has been of great interest.

Myosin II is a dimeric motor with two heavy chains and four light chains. Each myosin heavy chain is endowed with a pair of essential (ELC) and regulatory light chains (RLC). The heavy chain contains two main parts, i.e., the N-terminal globular motor domain and a long tail that participates in filament formation. As resolved in a crystal structure of the myosin head (Subfragment-1, S1), the motor domain contains the ATPase or active site, the actin-binding site, and a lever arm or light chain binding domain with ELC and RLC wrapped around it (1). The crystal structure of the human beta cardiac myosin motor domain has been recently resolved and shown to share common structural elements as found before (2). A lever arm amplifies the small conformational changes in the active site into a large displacement, whereby the actin filament is actively displaced by myosin. Light chains provide structural support to the otherwise floppy lever arm. The light chain-binding domain is followed by the heavy chain dimerization domain or Subfragement-2 (S2) region and continues as a tail that assembles into thick filament.

Myosins are mechanoenzymes that employ the energy from ATP hydrolysis to generate the force, driving actin filament movement and thereby muscle contraction. The biochemical actomyosin (AM) ATPase cycle proposed from earlier studies comprises different nucleotide states of the myosin active site, exhibiting strong and weak interactions with actin (3,4). In the absence of nucleotide, myosin associates with actin in a strong-bound ‘rigor’ state. ATP binding to the myosin active site dissociates the AM rigor complex. Myosin hydrolyzes the ATP to ADP and inorganic phosphate (Pi). In the dissociated/weakly-bound myosin state, the lever arm is reprimed in the pre-powerstroke configuration. The transition of AM from weak-to strongly-bound state and subsequent Pi release from the active site is associated with the generation of the first powerstroke, while ADP release with the second shorter powerstroke. During the pre-to post-powerstroke transition of the myosin head, lever arm rotates by about 70° during the 1^st^ powerstroke, and further following the ADP release. ADP release from myosin results in the formation of AM rigor state. The rigor state ends when the new ATP binds and the cycle continues. Despite following the same kinetic cycle, different myosin isoforms display large functional variability, i.e., distinct shortening velocities and force generation capabilities, owing to the differences in the transition rates among key AM states, allowing the motors to adapt to their physiological roles.

The *MYH7* gene located on chromosome 14 encodes β-myosin heavy chain (β-MyHC). The β-MyHC in complex with specific light chains, MLC1v and MLC2v, forms an active βM-II motor unit that is mainly responsible for driving the ventricular contraction in higher mammals. Apart from the left ventricle myocardium, the β-MyHC expression in slow-twitch skeletal muscle (M. Soleus) was established for humans (5,6) and other species, such as rats and rabbits (7,8). Along the same line, the β-MyHC with the hypertrophic cardiomyopathy (HCM) mutations were confirmed for the slow-twitch skeletal muscle M. soleus-derived myosin (SolM-II). Thus, in the patients with heterozygous HCM missense mutations, similar to cardiac samples, wild-type and mutant β-MyHC are expressed in skeletal muscle (M. Soleus). MyHC was largely accepted as a major functional determinant of the respective myosin complex. SolM-II with HCM mutations was perceived as an invaluable source to study the alterations in myosin properties which otherwise will be unavailable for research until or only if the patient undergoes myectomy or heart transplant. Since the skeletal muscle was relatively easily accessible, M. soleus-derived muscle fibers from HCM patients were employed to understand the impairment in muscle fibers mechanics (9-12). Besides, the purified motor proteins were analyzed in ensemble molecule measurements to assign the effects of specific HCM mutations in β-MyHC (5,13). While control experiments were performed to demonstrate the similar properties of myosin derived from soleus muscle and cardiac tissue, large variabilities were observed (13). The studies were performed in the context of the effect of mutations on the myosin function comparing soleus *vs*. cardiac mutant protein samples, where the ratio of mutant to wild type protein in different patient samples and even different human M. soleus-derived fibers likely affect the interpretations. The concrete evidence to substantiate the functional similarity between the myosin derived from cardiac and skeletal muscle under identical experimental conditions is still lacking. We, therefore, set out to perform systematic studies comparing myosin from rabbit M. soleus and left ventricular samples.

Contrary to the previous observations, we found significant differences in the ensemble-molecule actin filament gliding speed driven by motors extracted from the two-tissue sources. Using single-molecule analysis measurements, we further probed the specific chemo-mechanical features of the myosins for varied functional outcomes, i.e., gliding speed. Based on our studies, we conclude that regardless of the same myosin heavy chain isoform, the motors derived from different tissue sources exhibit diverse kinetic properties. The reason for this dissimilarity may become apparent when apart from a single major component, i.e., β-MyHC, the complete motor complex or holoenzyme is taken into account for the parallels. Our studies underpin the importance of studying the complete holoenzyme behavior. The accessory components and/or their modifications may attribute the key kinetic differences among motor complexes. This knowledge is crucial for future research aimed at unraveling the molecular mechanisms underlying individual motor isoform function and cardiomyopathies.

## Results

### Cardiac and slow skeletal myosin II-driven actin filament motility

Our main aim was to probe if the pure population of β-cardiac and slow skeletal myosin display comparable chemomechanical features as they share the same β-myosin heavy chain (β-MyHC) isoform. Note that full-length native β-cardiac and slow skeletal myosin are referred to as βM-II and SolM-II, respectively.

Native dimeric β-cardiac myosin motors were extracted from the rabbit heart ventricle and examined in the SDS-PAGE for purity. The heavy chain and light chain composition of the purified motors were examined for any contaminant atrial-specific α-cardiac myosins (Figure S1). The light chains corresponding to specific heavy chain isoforms were observed in the coomassie stained gels. No traces of either α-myosin heavy chain (α-MyHC) or essential (MLC1a) and regulatory light chains (MLC2a) were detectible in our samples, suggesting a pure population of βM-II. The distinct bands specific to β-MyHC and corresponding essential (MLC1v) and regulatory light chains (MLC2v) were observed. However, less than 5 % contaminant, which is beyond our detection limit, cannot be ruled out in our βM-II sample preparation. Similarly, slow myosin II were extracted from the rabbit type-I fibers generally found in slow-twitch muscles (M. soleus). The isolated proteins were scrutinized for their purity to avoid any contaminant fast skeletal myosin II isoform (PsoM-II), typically expressed in type-II fibers in M. psoas, as shown in Figure S2. The isolated motors were studied using actin filament gliding assay in our functional assays.

We measured ATP concentration dependence of the gliding velocities driven by βM-II and SolM-II. To avoid discrepancies related to the variation in the surface density of motor molecules, several ATP concentrations were tested in the same chamber (details in experimental procedure). As expected, the actin filament gliding velocity was sensitive to the ATP concentration for both βM-II and SolM-II. Figure 1 shows a plot of actin velocity as a function of ATP concentration from 5 µM to 2 mM ATP. The velocity increased with an increase in ATP concentration and followed the hyperbolic curve. To our surprise, the gliding velocities were almost threefold higher for βM-II than SolM-II at saturating ATP concentration. By fitting the distribution to Michaelis-Menten’s equation, the maximal velocities (*V*_max_) of 915 ± 35 nm/s and 281 ± 14 nm/s were estimated for βM-II and SolM-II, respectively. The *Km* was also about two-fold different, i.e., 35 µM and 16 µM for the βM-II and SolM-II, respectively.

**Figure 1.**
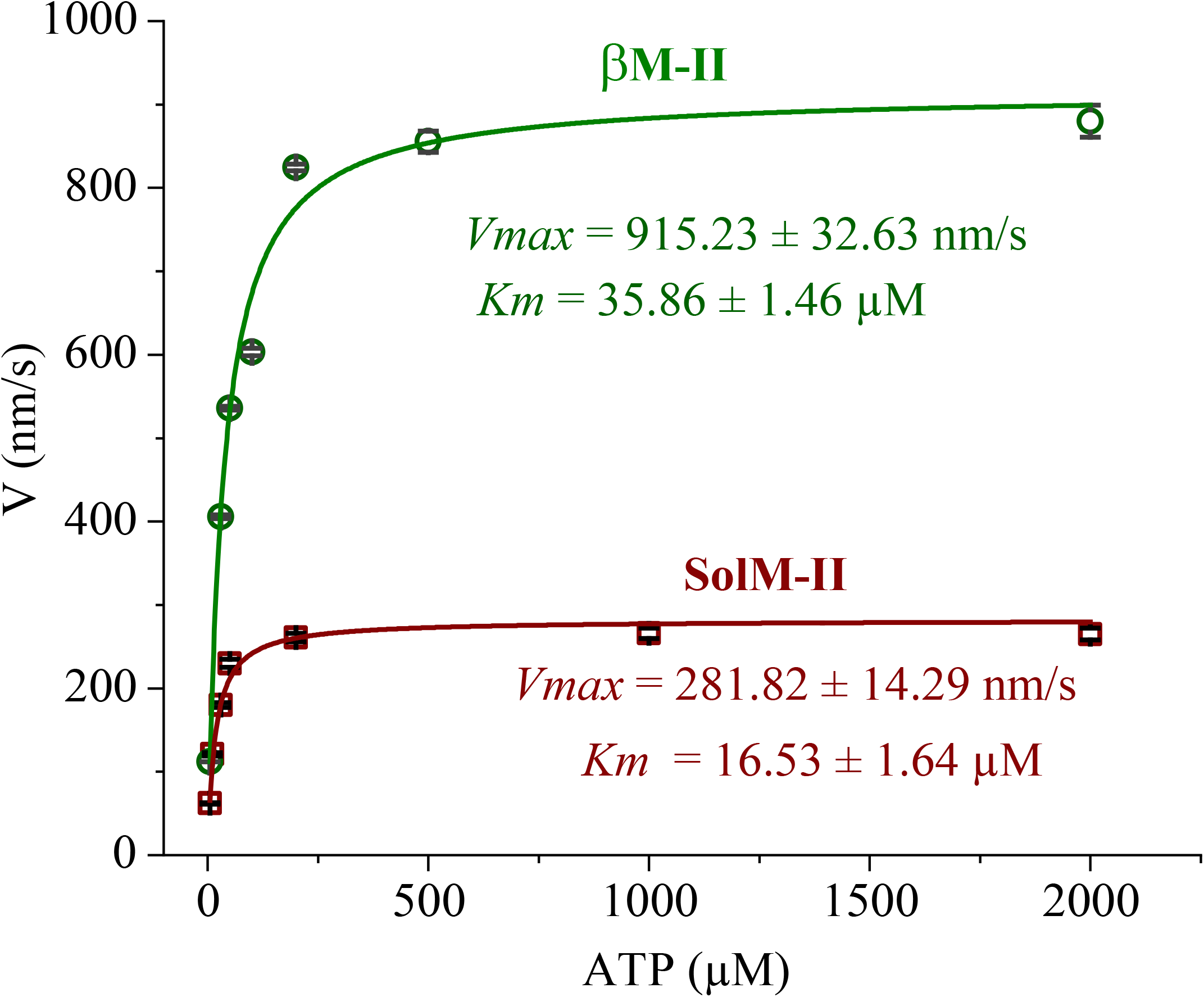
In vitro motility assay. Actin filaments gliding velocity on βM-II and SolM-II coated surface. The velocities were dependent on MgATP concentration for both motor forms and followed classic enzymatic reaction behavior defined by Michaelis Menten’s equation as a function of MgATP concentration. Each data point represent the average filament velocities ± SEM for corresponding ATP concentration. For each myosin, the data was fitted with Michaelis Menten’s constant yielding maximum velocity (*V*_*max*_) and ATP concentration required to achieve half-maximal velocity (*Km*). Gliding speed of 115-350 actin filaments were analyzed for each ATP condition. The experimental outcomes were reproducible with minimum three independent myosin preparations. SEM – standard error of mean.

Actin filament sliding velocities or unloaded shortening velocities are determined by the net rate of actomyosin cross-bridge cycling (14,15). The parameters that influence the sliding speed (*V*) include: (i) the lifetime of the strong AM bound state (*t*_on_), and (ii) the powerstroke size (*d*) of myosin, i.e., *V* = *d*/*t*_on_. During the actomyosin cross-bridge cycling, the fraction of the total ATPase cycle time myosin spends bound to actin is defined as the duty ratio. The duty ratio can change either by a change in the ADP release rate from the AM complex or the rate of weak-to-strong bound transition of AM (i.e., A.M. ADP.Pi to AM.ADP.Pi state). An additional factor that may influence ensemble-molecule sliding velocity measurements is the surface density of the myosin motors in the flow cell. The skeletal myosin II, a low duty ratio motor (< 0.05), showed motor density dependence of velocities (16,17). In our filament gliding measurements, similar myosin concentrations were infused into the flow cells. Therefore, the observed differences in the velocities for βM-II and SolM-II most likely resulted from altered powerstroke size (*d*) or the lifetime of the strongly bound state (*t*_on_) or both at saturating ATP concentration.

### Single molecule analysis of β-cardiac myosin

To gain the key details of the observed differences in the ensemble measurements, we employed single molecule analysis technique and investigated the biochemical and mechanical properties of individual motor molecules.

### AM detachment kinetics

The optical trapping based three-bead assay set up illustrated in Figure 2A is similar to previous reports (18,19). An actin filament is held tight between the two optically trapped beads. Native myosin motors are adsorbed onto the surface-immobilized nitrocellulose-coated glass bead. As depicted in Figure 2A, each bead position is tracked using quadrant detectors and recorded. The bead position records register the intermittent binding events between myosin (M) and actin (A), appearing as a reduction in the signal amplitude of free dumbbell noise/Brownian motion (Figure 2B). The mechanical interaction of myosin with actin is associated with the transition from ‘weak’ to ‘strong-bound’ states, concomitant with the generation of a powerstroke whereby the actin is displaced by the myosin molecule. Thus, for an individual actomyosin AM interaction, the duration of the reduced signal amplitude primarily indicates the steps following the first powerstroke. Note that since the strong bound AM.ADP.Pi state prior to the Pi release is short-lived i.e., <2 ms (20,21), it is expected to have minimal contribution to the lifetime of the observed interaction lifetime (t_on_). We assume that the AM binding event comprises the post-powerstroke strong bound ‘AM.ADP’ and ‘AM’ rigor states. The lifetime of the ‘AM’ rigor state depends on the ATP concentration, and increasing the ATP is expected to decrease the contribution of rigor states in the bound period. As the interaction lifetime (t_on_) becomes insensitive to the increase of ATP, the ADP release becomes rate limiting and thereby AM detachment rate is equivalent to rate of ADP release. The (AM) interaction events were measured at various ATP concentrations to study the ATP concentration dependence of the t_on_.

**Figure 2.**
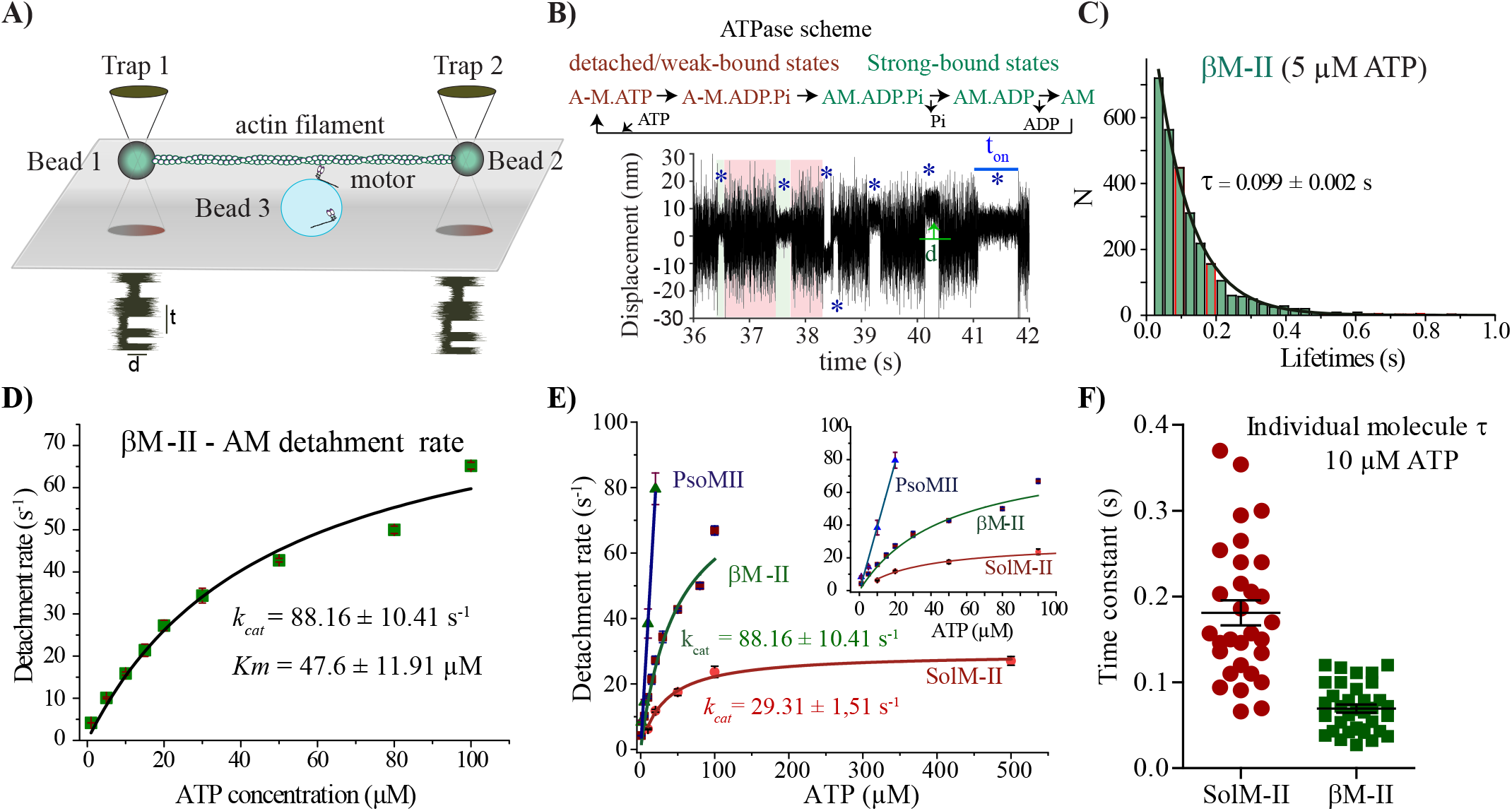
ATP concentration dependence of AM bound lifetimes for βM-II and SolM-II. **(A)** Optical trap setup for three-bead assay. Note that the various components of the setup are not drawn to scale. **B)** Original data trace displaying the bead position signal over time. Example shows single myosin molecule interacting with an actin filament at 5 µM ATP. Several AM interaction events are indicated with blue asterix. t_on_ is the duration of AM association. The green arrow illustrates the displacement (d) from average unbound to bound position. The AM bound and unbound states are depicted in simplified ATPase scheme and corresponding signal change is shown with green and red background on the data trace, respectively. **C)** AM interaction events t_on_ measured at 5 µM ATP are plotted in a histogram. The average lifetimes (τ) were calculated by least-squares fitting of a histogram with single exponential decay function. **D)** The AM detachment rate is the inverse of AM-bound average lifetime (τ). Detachment rates (1/τ) at increasing concentration of ATP from 1 to 100 μM derived from τ as shown in (C) are plotted for βM-II. ATP concentration dependence of the detachment rate follows the Michaelis-Menten kinetics and thus fitted with the function, *v* = *V*_*max*_*x/(*K*_*m*_+x). Maximum detachment rate (*V*_*max*_ or *k*_*cat*_) and ATP concentration at ½ *V*_*max*_, i.e., *Km* was derived, R^2^ = 0.97. **(E)** ATP concentration dependence of the AM detachment rate (1/τ) as a function of ATP concentration are compared for βM-II, SolM-II and PsoM-II. Inset shows the clear kinetic difference observed at lower ATP concentrations among the three-myosin forms. Rates for βM-II and SolM-II fitted with Michaelis-Menten function, and for PsoM-II, with linear regression. For PsoM-II, kT from linear regression was 3.96 ± 0.31 μM^-1^ s^-1^; for βM-II, *k*_*cat*_ = 88.16 ± 10 s^-1^, *Km* = 47.6 ± 11 μM, for SolM-II, *kcat* = 29.3 ± 1.5 s^-1^, *Km* = 31.6 ± 3.7 μM. **F)** Scatter plot depicts the time constants determined from measurements at 10 µM ATP from individual molecules for SolM-II (N = 30) and βM-II (N = 35), and are significantly different P< 0.0001 (two tailed t-test). The error bars are average ± standard error of mean (SEM). Altogether, 231 individual βM-II molecules and 36000 AM binding events were identified and analyzed. At 1 μM ATP, N = 59, n = 8211; 5 μM ATP, N = 18, n = 3219; 10 μM ATP, N = 48, n = 7240; 15 μM ATP, N = 21, n = 2699; at 20 μM ATP, N = 22, n = 3893; 30 μM ATP, N = 10, n = 1050; at 50 μM ATP, N = 63, n = 5375; 80 μM ATP, N = 14, n = 4659; at 100 μM ATP, N = 15, n = 1381. The single-molecule experiments were performed with myosins from at least three separate βM-II preparations. Note that lifetime measurements at 50, 80 and 100 µM ATP were possible with the fast triangular wave of 600 Hz on one of the beads so that the binding events are discernible. For SolM-II, 10 µM ATP N = 30, n = 4344; 20 µM ATP N = 53, n = 7336; 50 µM ATP, N = 16, n = 1651; 100 µM ATP, N = 40, n = 4937; 500 µM ATP, N = 40, n = 2099. For PsoM-II, in total 30 individual myosin molecules were measured for 1, 5, 10, and 20 μM ATP conditions. N = number of individual myosin molecules and n = number of AM association events. The event lifetimes for βM-II and SolM-II were compared between different ATP concentrations using the nonparametric Mann-Whitney U test, which yielded the statistical differences in t_on_ with P <0.0001.

We used the varying concentrations of ATP from 1 to 100 µM and measured the duration of AM bound state (t_on_) at defined ATP concentrations. Average t_on_ (τ) for each ATP concentration is determined, an example as shown for 5 µM ATP in Figure 2C. The reciprocal of τ is used to calculate the AM detachment rate (1/t_on_). As shown in Figure 2D, the ATP concentration dependence of the detachment rate follows the Michaelis-Menten kinetics, yielding maximum detachment rate of 88 s^-1^ and the Michaelis constant, *Km* i.e., the ATP concentration to achieve half maximal detachment rate was estimated to be about 48 µM. Note that the measurements above 100 µM ATP were not feasible, as the interaction events were too short for reliable detection. In Figure 2E, the detachment rates of cardiac myosin βM-II were compared with the slow myosin SolM-II and fast PsoM-II. SolM-II displayed nearly 3-fold lower maximum detachment rate of ∼ 30 s^-1^ than βM-II. *Km* was also lower for SolM-II, i.e., ∼31 µM. βM-II exhibited the AM detachment kinetic properties intermediate between the fast PsoM-II and slow SolM-II. The detachment rates for PsoM-II was not possible to determine as it’s reported to be in the range of 500 s^-1^ (22), and thus beyond the detection limit of our current optical trap set up.

Thus, the observed differences in ensemble actin filament gliding velocities were recapitulated in the AM dissociation rates at single-molecule level for SolM-II and βM-II. The *Km* estimated from the motility experiments were although lower but showed similar trend to the detachment kinetics in single-molecule experiments, thus reinforcing that the requirements for the ATP concentration to reach half-maximal velocity is shifted to higher values for βM-II.

We also compared the average lifetimes derived from individual βM-II and SolM-II molecules to probe 1) if the heterogeneous population of the motor molecules causes the observed difference in the kinetic rates and 2) if a cluster of motor molecules with identical kinetic properties is apparent (Figure 2F). While larger variation was observed for SolM-II, a subpopulation of motors with similar kinetic properties between SolM-II and βM-II could not be distinguished. Altogether, βM-II and SolM-II revealed significantly different kinetics of actomyosin detachment.

### β-cardiac myosin powerstroke size

Apart from AM detachment kinetics, the stroke size can influence the motor-driven speed of actin filaments. To estimate the stroke size of native βM-II, the AM binding events were analyzed for the displacement from mean dumbbell position. The individual event displacements were plotted and a shift-of-histogram method was employed to determine the mean displacement of the myosin motor as shown in Figure 3A. AM binding events measured at 5 µM ATP concentration yielded the average stroke size of 5.32 nm for βM-II. Consistent with our previous studies (23), we measured total stroke size of about 6 nm for SolM-II (23).

**Figure 3.**
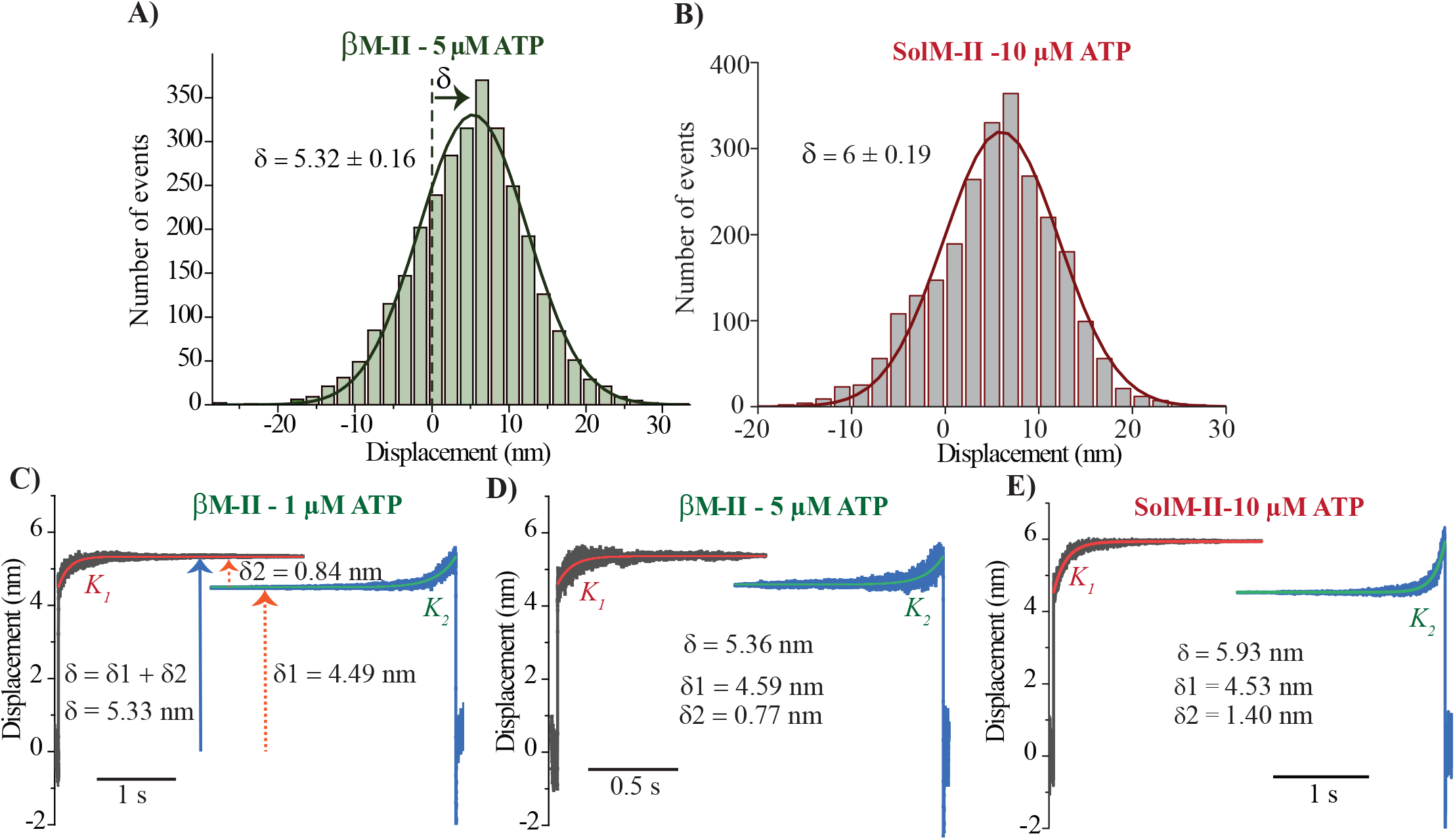
Powerstroke size of β-cardiac myosin. **A and B)** The histogram with displacement events measured at 5 µM ATP for βMII and at 10 µM ATP for SolM-II. The average stroke size was estimated by using shift-of-histogram method (68). Least-squares fitting of event distribution with Gaussian function yields average powerstroke, δ = 5.32 ± 0.16 nm and 6 ± 0.19 nm for βM-II and SolM-II, respectively. **B -D)** Ensemble averaging of the individual AM association events. The beginning and the end of the several individual AM binding events were synchronized and fitted with single exponential functions to estimate the substeps. The beginning is shown as a forward fit (red points) and the end (green points) as a backward fit. Average stroke size (δ), and displacement size corresponding to the 1^st^ and 2^nd^ stroke is indicated as δ1 and δ2, respectively. K_1_ - reaction rate for the ADP dissociation and *K*_*2*_ - ATP induced AM dissociation at respective ATP concentration. *K*_*2*_ = 4 S^-1^ and 10.2 S^-1^ at 1 and 5 µM ATP, respectively. For 1 µM ATP, N = 16, n = 987; for 5 µM ATP, N = 7, n = 500. For SolM-II at 10 µM ATP, N = 11, n = 520. N = Number of molecule, n = number of events. Important to note that for ensemble averaging, the AM attachment events with a lifetime of minimum 0.05 s or longer were selected. The method is described in detail in Veigel et al.(24) and Blackwell et al. (25)

The total stroke size is expected to contain substeps that are associated with the Pi and ADP release, as the ATP undergoes ATPase cycle. The 1^st^ powerstroke, linked to the Pi release, is the beginning of the mechanical interaction between actin and myosin. Within the AM bound period, the 2^nd^ powerstroke associated with the ADP release should take place as the myosin transitions from AM. ADP to AM state. Since the overall size of the steps is small, it is difficult to extract the substeps from normal displacement over time records, as the amplitude of Brownian motion of the trapped bead is larger compared to the stroke size. Moreover, as the 1^st^ powerstroke occurs in less than 2 ms after binding to actin filament, damping effect of the trapped bead masks the myosin’s conformational change following the initial AM attachment. To resolve the substeps Veigel et al. (24) introduced the ensemble-averaging method, whereby synchronizing sufficiently long events at the start and the end of the events were used to extract the hidden information in the noise amplitudes with high precision. We employed this method to separate the substeps, i.e., 1^st^ and 2^nd^ powerstroke. Recently reported matlab-based program, Software for Precise Analysis of Single Molecules (SPASM) (25) was used to synchronize the events at the beginning and end of the event as shown in Figure 3C-E. At lower ATP concentrations, the bound durations of AM are expected to contain sufficiently long time traces to derive the amplitude of 2^nd^ stroke (i.e., transition from AM.ADP to AM). For βM-II, the analysis for two different ATP concentrations (1 and 5 µM) revealed the total stroke size and amplitude of each step to be ∼4.5 nm and ∼0.8 nm for the 1^st^ and the 2^nd^ stroke, respectively. We also compared the substeps measurement with the SolM-II, which was 4.53 and 1.4 nm, respectively. Interestingly, the analysis revealed similar amplitude for 1^st^ powerstroke of about 4.5 nm for both βM-II and SolM-II. We also determined *K*_2,_ the transition rate from AM rigor to ATP bound M state or detached state, which as expected increased with increasing ATP concentration, i.e., 4 s^-1^ at 1 µM and 10 s^-1^ at 5 µM, respectively (Figure 3C-D). *K*_1_, which indicates the transition from AM.ADP to AM state or the rate of ADP release is however difficult to determine reliably from this analysis, as the positive feedback applied to improve the signal to noise ratio interferes with the initial attachment amplitude signal. Nevertheless, the ADP release rate was reliably estimated from the ATP concentration dependence of the AM association lifetimes as shown in Figure 2D and E. Thus, while the first powerstroke size was similar between βM-II and SolM-II, a small difference in especially the 2^nd^ stroke was observed.

### β-cardiac myosin stiffness

The rigidity of the motors is key to their force-generating ability as the motor interacts with the filament and undergoes nucleotide-dependent conformational changes. Variance-covariance analysis (26,27) was employed to estimate the stiffness of βM-II. Extent of the noise amplitude reduction of both the trapped beads during AM interaction was used to derive the stiffness of the interacting motor head. Our recent study and other reports (23,28,29) have established that a single motor head interacts with the actin filament during single association-dissociation events. In Figure 4A, stiffness measurements performed at two different ATP concentrations, i.e., 1 and 50 µM ATP are shown. We estimated high cross-bridge stiffness of about 1.83 ± 0.09 pN/nm at 1 µM ATP, while it reduced to about 0.88 ± 0.08 pN/nm at 50 µM ATP. We demonstrated recently for slow SolM-II, that the ADP bound AM state has lower stiffness than the AM rigor state (23). For SolM-II, stiffness of 1.75 ± 0.1 pN/nm and 0.5 ± 0.04 pN/nm was determined at 10 and 500 µM ATP, respectively (Figure 4B). It is conceivable that at low ATP concentration (e.g., 1 µM), within the lifetimes of AM interaction, the periods of reduced noise primarily comprises the ‘rigor’ states, whereas the duration of ADP bound AM states constitutes a relatively minor fraction. Therefore, the time-averaged stiffness is dominated by the rigor cross-bridge state. At high ATP, however, the lifetime of ‘rigor’ state is reduced and the duration of AM bound ADP state forms a substantial fraction of the total bound duration. Thereby, the time-averaged stiffness primarily represents the ADP state or the average of the two states (‘AM.ADP’ and ‘rigor’) depending on each state’s contribution. Figure S3 shows ATP concentration-dependence of the measured stiffness where the shift towards lower stiffness values is evident with increased ATP, corresponding to average of ADP-bound AM state. From our analysis of maximal detachment rates (i.e., 88 s^-1^, which means the average event lifetime of ∼11 ms), ATP concentrations of more than 100 µM or higher would be required to acquire predominantly ADP bound AM states. For βM-II, the measurements above 50 µM ATP resulting in shorter event lifetimes were insufficient for reliable cross-bridge stiffness estimation with either variance-covariance or triangular wave method.

**Figure 4.**
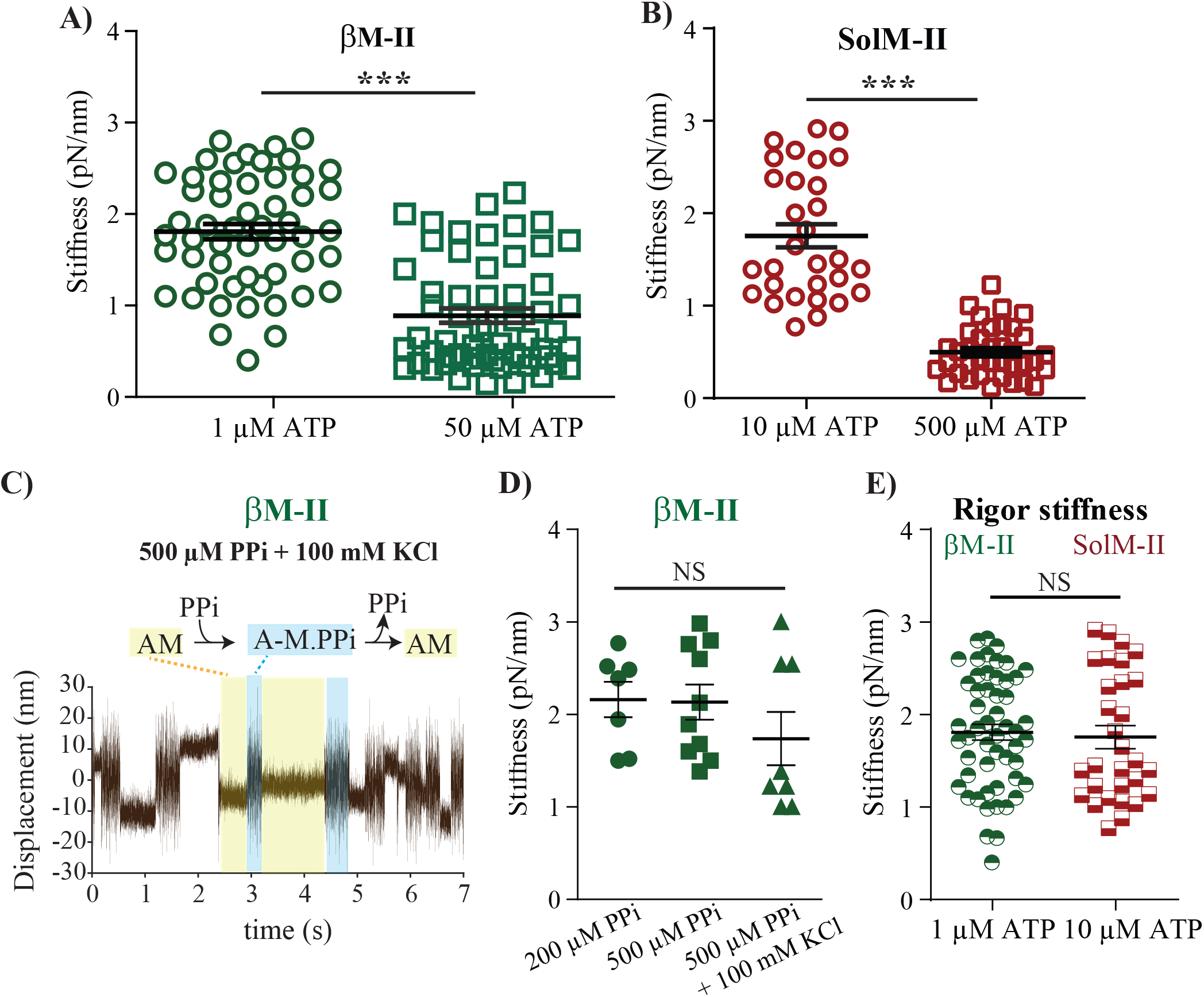
Stiffness of β-cardiac myosin. measured at high and low ATP concentrations. **(A**) Scatter plot with mean and SEM shows the stiffness measured for β-myosin and at 1 and 50 µM ATP concentrations. Each data point displays the stiffness measured from an individual molecule. β-myosin at 1 μM ATP (N = 54) and 50 μM ATP (N = 59) with the average stiffness of 1.83 ± 0.09 pN/nm vs 0.88 ± 0.08 pN/nm, respectively, with P < 0.0001, revealing highly significant difference. **(B)** Scatter plot shows stiffness at 10 μM and 500 μM ATP for SolM-II. For SolM-II at 10 μM ATP (N = 30) and 500 μM ATP (N = 37), P < 0.0001. Rigor stiffness between the β-myosin (at 1 µM ATP) and SolM-II (at 10 µM ATP) was not significantly different, P = 0.66. Independent sample t test was used to calculate the statistical significance. **C)** The AM cross-bridge stiffness measured in the presence of pyrophosphate (PPi). The stiffness for three different PPi conditions was comparable and not significantly different; P < 0.05 with Bonferroni’s multiple comparison test. 200 µM PPi; N =7, n= 576, 500 µM PPi; N = 11, n= 831; 500 µM PPi+100 mM KCl; N = 6, n= 716. N = number of individual motor molecules, n= number of binding events.

Nevertheless, we observed similar trend for βM-II as the SolM-II that the rigor stiffness is higher and increased ATP concentrations lead to lower stiffness values, suggesting that the AM.ADP state possesses at least two-fold lower AM cross-bridge rigidity.

### Rigor stiffness in the presence of pyrophosphate PPi

To further confirm that the state with a higher stiffness represents a rigor state, we mimicked the strong-bound rigor-like AM state and measured the stiffness. We acquired cues from previous studies, where pyrophosphate (PPi) was used to attain primarily the near-rigor state (30). Binding rate of M.PPi to actin from solution studies is estimated to be 10^7^M^-1^s^-1^(31). No working stroke was observed in the presence of PPi, leading to the conclusion that the lever arm position remains similar to that of the rigor state (30). In our analysis, the data records showed clearly defined rapid rebinding events in the presence of PPi (Figure 4C). The ionic strength was raised to decrease the binding and detect individual interaction events. In principle, PPi was used in the reaction mixture to promote unbinding of the rigor complex, and high affinity of the PPi enabled collection of individual binding/unbinding events. It was feasible to acquire a large number of events from individual motor molecules. The binding events were analyzed and the stiffness was derived for single myosin molecules using variance-covariance method. As shown in Figure 4D, high stiffness of about 2 pN/nm was noted for different pyrophosphate conditions, which was similar to the values obtained at lower ATP concentrations, i.e., 1 and 10 µM ATP for βM-II and SolM-II, respectively. Thus, we interpret that stiffness of actomyosin cross-bridge measured at low ATP represents the rigor stiffness, which was found similar for both βM-II and the SolM-II as shown in Figure 4E.

### Average stiffness vs. lifetimes

We observed large variability in the time constants (τ) from individual molecules as seen in Figure 2F. We therefore wondered if there is a direct correlation between the lifetime and the measured stiffness for individual molecules i.e., whether longer average lifetime yields high stiffness for the specific myosin head and the shorter τ corresponds to the lower stiffness. In Figure 5, the plots of stiffness over time constants for individual molecules of βM-II and the SolM-II are shown. We found no obvious link between the time constants and the measured stiffness for single molecules.

**Figure 5.**
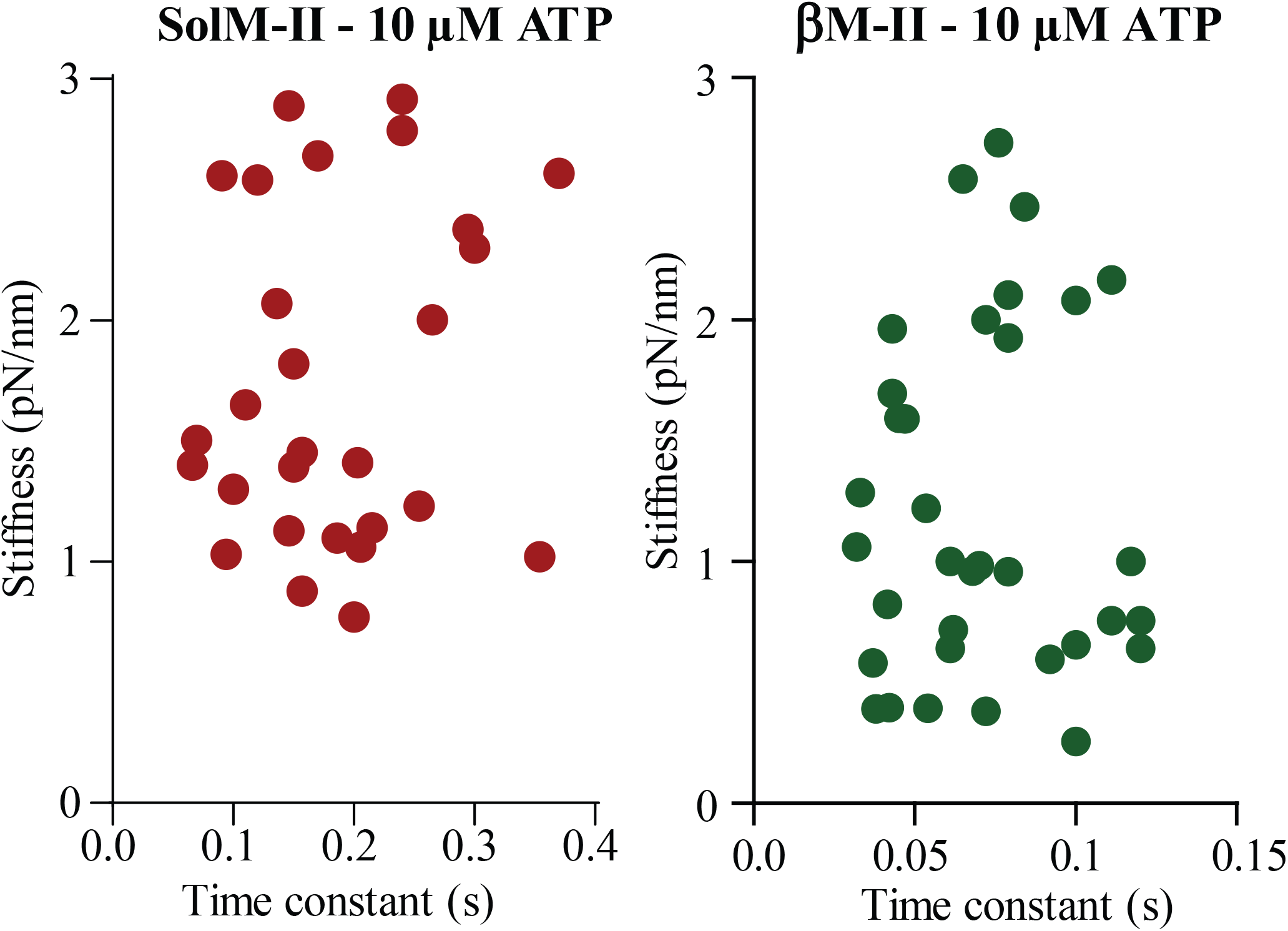
Stiffness vs average lifetimes of AM-bound states. For individual molecules, the interaction events were analyzed for time constants (τ) and the cross-bridge stiffness. τ is plotted against the measured stiffness for each molecule to check if there is any correlation between the average duration of interaction and measured stiffness. The lifetimes and stiffness measured for individual molecule are compared for the AM interaction events collected at 10 µM ATP. βM-II, N = 34 and SolM-II - N= 30.

## Discussion

The *MyH7* gene product β-MyHC is the predominant myosin heavy chain isoform expressed in the ventricular myocardium and aerobic slow-twitch skeletal muscle fibers. In previous studies, the myosin heavy chain isoforms expressed in the distinct muscle fibers were shown to be the major determinant of the contractile velocity of the respective muscle fibers. Therefore, slow myosins isolated from M. soleus were considered an alternative to examine cardiac muscle physiology and pathophysiology, such as in human cardiomyopathies. In the present study, we investigated the precise kinetic and mechanical features of the βM-II and SolM-II motors derived from ventricle and slow skeletal muscles using ensemble- and single-molecule assays (summarized in Table 1). Actin filament sliding velocities were found significantly lower for SolM-II than βM-II. The differences in the velocities are in agreement with the very recent work for the two motor complexes (32). The ATP concentration dependence of the velocity (*Km*) was shifted toward higher concentrations for cardiac myosins. Single-molecule optical trapping measurements assigned the reason behind the slower velocities to the slower dissociation kinetics of the ADP for SolM-II. The mechanical parameters such as powerstroke size and the motor stiffness were comparable. Remarkably, the recently found feature for SolM-II (23), i.e., the change of crossbridge stiffness during actomyosin ATPase cycle, was ascertained for the βM-II as well. Accordingly, AM.ADP state displayed lower stiffness than the rigor stiffness for βM-II. Collectively, the functional differences identified in these studies recommend careful considerations while using skeletal muscle myosin or fibers as a replacement to study cardiac muscle pathophysiology.

**Table 1.**
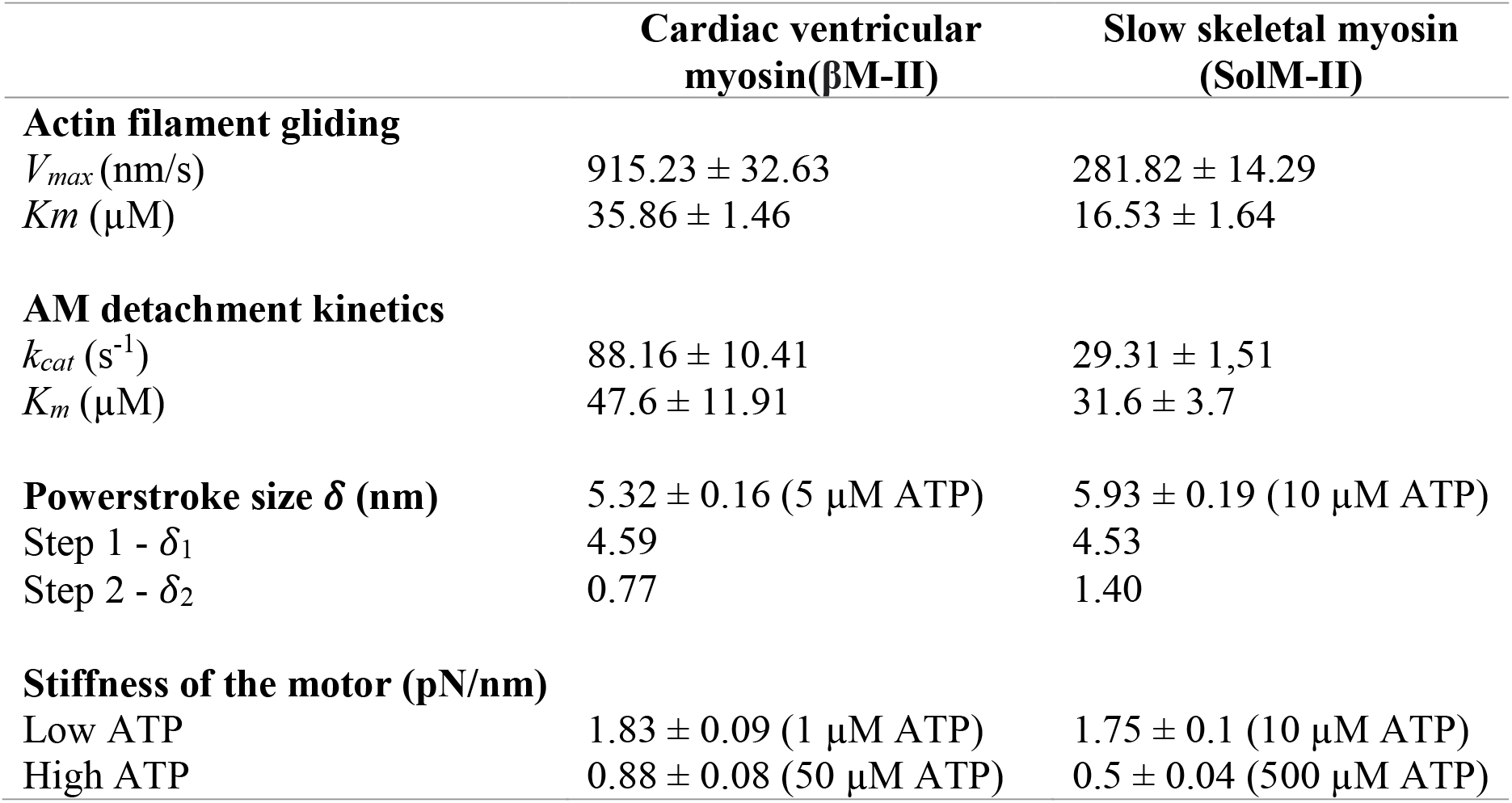
Summary of parameter derived from ensemble molecule and single-molecule studies.

### Relation to earlier studies on soleus and cardiac myosin

The actin filament gliding velocity of the βM-II of about 1 µM/s is comparable to the previous reports for rabbit, human and porcine ventricular myosin (32,33). Our observed maximum AM detachment rate of 88 s^-1^ for full-length rabbit β-cardiac myosin is slightly faster than the previous reports. The rate of ADP release measured for expressed human single headed beta cardiac myosin was 64 s^-1^ (34), while single-molecule optical trapping of porcine cardiac myosin estimated detachment rate of 74 s^-1^ (35). Species-specific subtle variations or the experimental conditions, e.g., temperature, single *vs* double headed myosin, myosin complex with all the subunits or without RLC employed in the measurements are expected to have an influence on this parameter. Nonetheless, similar values for rate of AM detachment and ADP release suggest that the structural transition corresponds to the release of ADP.

Our *Km* values from the single-molecule studies are slightly higher than those for motility experiments i.e., the ATP concentration dependence higher to reach half-maximal *Vmax* (*Km* of 50 and 35 µM for cardiac and 31 and 16 µM for slow skeletal myosins, respectively). One possible explanation for the difference in apparent *Km* values in the filament velocity is assisting load by the cycling AM cross-bridges - albeit low - in the mechanical sliding of actin filaments causing the faster detachment and availability of nucleotide-free myosin heads. In other words, detachment is faster in the motility experiment configuration in comparison to the single-molecule measurements. Conversely, in single-molecule trapping conditions, individual myosin heads operate independently. Besides, the surface immobilization of the myosin molecules on different surfaces, i.e., nitrocellulose and BSA coated surface for single-molecule and gliding assay, respectively, may impact the AM detachment kinetics and the gliding speed. The *Km* value of 35 µM from motility assays is in close agreement with previous measurements for rabbit and rat cardiac myosins of 25-40 µM and 43 µM, respectively (36,37). The small differences may be explained by the differences in the assay conditions, i.e., motility of the myosin-coated beads along the actin cables (37) and a species-specific difference (36). For the powerstroke size measurements, similar to previous observations, two substeps with average displacements of about 4.5 and 0.8 nm were noted. Our estimated total stroke size of 5.3 nm for βM-II is lower than the previous single-molecule studies using porcine ventricular myosin (6.8 nm) (35). For rat ventricular myosin, the values between 3-8 nm were estimated in *in situ* studies (38). The authors also indicated load dependence of the stroke size, i.e., higher load producing shorter displacement of 3 nm and lower load allowing stroke size of 8 nm. Again, the experimental conditions and species-specific differences likely contributed to the observed small variabilities.

### AM cross-bridge stiffness

The change of AM cross-bridge stiffness during the ATPase cycle is rather a very recent observation (23). Interestingly, for βM-II we made similar observation. With increasing ATP concentration, the trend toward lower stiffness, i.e., from 1.83 pN/nm to 0.88 pN/nm was observed (Figure 4A and S3). We assign these two stiffness values to the ‘rigor’ and ‘ADP-bound’ cross-bridge states, respectively. Although due to limited time resolution, mainly AM-ADP states could not be achieved, the data points towards two different cross-bridge stiffnesses as the myosin head undergoes the cross-bridge cycle. We previously predicted that change of stiffness within AM interaction time may be a common feature among different isoforms of motors and that measured value for stiffness will depend on the duration of the respective state. Our observation for βM-II further strengthens this proposal. Besides, using PPi we could directly probe the rigor stiffness for the motor complexes, which was found comparable for the SolM-II and βM-II. Slightly higher average stiffness values in PPi experiments compared to those observed at low ATP concentration may suggest a small contribution from ADP-bound states of AM cross-bridges.

Mechanical work done *(W)* per ATP hydrolyzed is estimated previously from muscle fiber and single-molecule studies (39-41). The estimated values for *W* were 30 pN nm (39), 27 pN nm (40) from muscle fiber mechanics, and 11 pN nm per ATP from single-molecule studies (41). Using our powerstroke size and stiffness values, we could utilize a function for the potential energy of a spring, i.e., 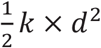 (*d* - overall stroke size/displacement, *k –stiffness*/spring constant) to calculate the mechanical work done per ATP molecule per AM crossbridge cycle. Using overall stroke size of 5.3 and 6 nm and stiffness values of 1.83 and 1.75 pN/nm for βM-II and SolM-II, respectively, the work done were calculated to be 25.7 and 31.5 pN nm. These estimates are in close agreement with the estimates from fiber studies. Note that here we used the rigor stiffness of the AM crossbridges, i.e., the state when the powerstroke is completed.

### Possible source of differences in the β cardiac and M. Soleus M-II

Diverse MyHC isoforms are endowed with specific ELC isoforms, for example, the slow myosin ELCs are distinct from the ones that associate with fast myosins. For both fast and slow skeletal myosin isoforms, two essential light chains with size ranging from 17–27 kDa have been identified. For fast myosin, MLC1f and MLC3f (42), and for slow myosin MLC1sa and MLC1sb/v ELCs are known (43). In vivo, the two ELC isoforms can thus assemble with MyHCs to generate three different forms of the myosin hexamer with respect to ELCs, comprising either a homo- or heterodimer. In earlier work, the significance of the ELCs was examined using reconstitution approach. Fast, slow skeletal and cardiac myosin ELCs were swapped to produce different hybrid single-headed myosins. Biochemical analysis of these hybrid motors revealed deviation of actin-activated ATPase kinetics from that of naturally existing combination of myosin complexes, corresponding to the change of ELCs (44). In subsequent studies, non-native ELC isoforms exchanged into the permeabilized muscle fibers showed ELC isoform-dependent effect on the maximal shortening velocity (45). The two slow ELC isoforms MLC1sa and MLC1sb reconstituted in the fast M. psoas fibers showed greater reduction in the shortening velocity compared to the fast ELC isoforms. However, the difference between the two isoforms MLC1sa and MLC1sb was not significantly different. An effect of the ELCs was also investigated using an in vitro motility assay (46). Here, the long MLC1f and short MLC3f isoforms of ELCs were compared. MLC3f containing heavy meromyosin (HMM) moved actin filaments approximately twice as fast as MLC1f -containing HMM. The study revealed that the shorter ELC promotes the faster gliding speed.

In the current study, the obvious difference in the composition of the two holoenzymes is an extra ELC, i.e., MLC1sa in myosin originating from M. soleus. Based on past reports, it appears likely that the kinetic features observed for SolM-II are due to the existence of a longer ELC, MLC1sa (27 kDa) while the βM-II is equipped with MLC1sb/v (24 kDa). By comparing individual molecule attachment lifetimes (t_on_), we probed if the long ELC, i.e., MLC1sa is responsible for the slower AM detachment kinetics of SolM-II. We examined if the population corresponding to the similar composition as βM-II is discernible in our single-molecule analysis. However, the time constants from individual SolM-II molecules did not reveal two distinct populations (Figure 2F), corresponding to an effect of a shorter or longer ELC on the detachment kinetics. In a dimeric motor, the existence of a population of hetero- and homodimers of motors, and the type of individual trimeric complex’s (β-MyHC-MLC1sa-MLC2v or β-MyHC-MLC1sb-MLC2v) affinity and activity is likely to determine the measured time constants for an individual molecule. Besides, in a dimeric form, even if a single head interacts with the actin filament, the non-interacting head is known to have influence on its activity. One question arises how ELC influences the actomyosin crossbridge properties.

Several studies support the viewpoint that the long N-terminus of ELC directly interacts with actin (47-51). About 40 aa N-terminal extensions in long ELCs are rich in lysine and proline residues. A common view is that the weak interactions between the positively charged N-terminus of the ELC and negatively charged C-terminus of actin modulate the kinetics of AM cross-bridge cycling, reducing the AM complex detachment and consequently filament sliding velocity. In other work, however, contradictory results were observed. Instead of single-headed myosin (S1), when more physiologically relevant motor complexes such as full-length or dimeric myosin HMM were employed, the N-terminal region of long MLC1 reduced the affinity of myosin for actin, rather than strengthening (52).

Previous studies, including our own, highlighted that the variants of light chains critically influence motor function (22,53). We have demonstrated it for the myosin light chain 2 (RLC). Accordingly, despite its position distal to the main motor domain, MLC2v decelerated the fast skeletal myosin-driven motility, implying an allosteric effect on the ATPase cycle. Based on our observation with the RLC’s impact on the AM dissociation rates and actin filament gliding speed, it appears likely that the reduction in the velocity may be a direct consequence of the effect of long ELC isoform on the ATPase kinetics, e.g., on the ADP release. The FRET and Cryo-EM experiments suggested that the N-terminus of ELC may interact with the SH3 domain (Src-homology domain 3, comprising a six-stranded β barrel) of the globular head (54). This interaction is assumed to be a bridge between the ELC and actin filament to encompass about 8 nm distance. Conceivably, through its interaction with the N-terminal SH3 domain of MyHC, ELC may be responsible for modulating the steps of the ATPase cycle, such as ADP release. At saturating ATP concentration, the slower ADP release causing slower AM detachment can directly affect the moving filament. The differences in the fraction of the AM crossbridges in the post-1^st^ powerstroke state (i.e., ADP state) are likely to translate into the different velocities.

Although many studies support the proposal that the ELC linking to the actin filament is possible, it remains to be seen whether this feature is directly responsible for the reduced velocities for the motor forms with long ELC’s N-terminal link. In this state, the additional weak interaction between actin and myosin via ELC should be overcome following the ADP release and consequent 2^nd^ powerstroke, allowing AM complex to dissociate after ATP binding to the active site. The two proposals, i.e., direct or allosteric effect on ATPase cycle and braking effect through engaging with actin filament, however, may not be entirely exclusive. One possibility is that the ELC N-terminal association to actin may slow down the ADP dissociation if it imposes a resisting load on the myosin head. This effect perhaps is difficult to notice at the single-molecule level but possible in the ensemble-molecule experiments such as actin gliding assay, where several motors are simultaneously engaged to drive actin movement. In the light of contradictory results from previous studies concerning the long ELCs role to strengthen or weaken the AM interaction, additional specific experiments in the future would be required to verify these suggestions and to examine ELC’s modulatory role.

### RLC phosphorylation

β-cardiac myosin RLC for several species have shown to be nearly 40 % phosphorylated (55,56). The RLC phosphorylation is indicated to aid the movement of myosin heads away from the thick filament backbone from so called ‘OFF’ to ‘ON’ state (57-59), increasing the fraction of disordered myosin heads that are readily available for interaction with the actin filament. Thus, RLC phosphorylation is expected to increase the fraction of myosin heads for force generation. Consistent with this proposal, phosphorylation of cardiac RLC is shown to increase the isometric force production (56,59). For slow-twitch muscles, however only a small fraction of about 8 % is found to be phosphorylated (60). In vivo, the myosin molecules are closely spaced/parallel to the thick filament backbone and requires to be lifted off the backbone to make contact with the thin filament. The RLC phosphorylation is understood to play a major role in this mode of myosin ‘ON’ conformation. In *in vitro* experiments, the myosin heads are expected to immobilize in the upright orientation to support its association and displacement of actin filaments. The extent of impact phosphorylation may display in this experimental set up is rather unclear. However, subtle effects of RLC phosphorylation on the velocities were observed in in vitro motility experiments (61,62). For the purified motors employed in our studies, we did not find significant difference in the phosphorylation levels of the βM-II or SolM-II. Our observation was consistent with a recent report on RLC phosphorylation levels (32). Likely reason is that the phosphorylation is lost during the protein extraction. Overall, nearly threefold difference in the gliding speed seen between βM-II and SolM-II is less likely to be caused by variable RLC phosphorylation levels of the motor complexes.

In conclusion, these observations indicate that the MyHC alone does not determine the functional outcome of the motors but the overall composition of the hexameric complex and the interaction between individual components of the holoenzyme modulate the chemo-mechanical coupling to serve the physiological role. AM detachment kinetics, which is limited by the ADP release rate, found to be the determining factor guiding the function of the motors when they are required to perform as a cohort. It appears that, with a small variation in their composition, the two motors are adapted to operate differently in their respective physiological milieu, i.e., β cardiac myosin motors are engaged in generating maximal force during systole to pump blood from the ventricle, whereas slow skeletal myosin can sustain the load under voluntary signal.

The current study has implications in our understanding of the molecular basis of various tissue-or organ-specific roles of the myosin isoforms. Besides, it highlights the significance of the in-depth investigation when the corresponding motor complexes are employed to understand the physiology and pathophysiology associated with motor components. While the current studies primarily refer to the rabbit myosins, it raises a possibility that the effects of the cardiomyopathy mutations in SolM-II backbone may vary from the ones in βM-II. Further studies with human tissue-derived motors will, however, be required to ascertain the choice of substitute proteins for important disease-related examinations.

### Experimental procedure

#### Native myosin II

Full length cardiac myosin II (βM-II) and slow myosin (SolM-II) were isolated from rabbit ventricular tissue and *M. soleus* muscle in the high salt extraction buffer (0.5 M NaCl, 50 mM HEPES, pH, 7.0, 5 mM MgCl_2_, 2.5 mM MgATP, and 1 mM DTT) as previously described (63,64). The cardiac myosin was isolated from 50-100 mg ventricular tissue powder for each experiment as described in (65) with some modifications. Isolated myosin was aliquoted, flash frozen in liquid nitrogen and stored at -80°C in 50% glycerol.

The muscle and cardiac tissues were collected from the New Zealand white rabbits, Crl:KBL (NZW). The animals were euthanized as per the guidelines from German animal protection act §7 (sacrifice for scientific purposes). In this study, we used shared organs originating from the animals approved for experiments with authorization number 18A255. The animals registered under reference number G43290, were obtained from Charles River France. All the procedures were carried out in accordance with relevant guidelines and regulations from the Lower Saxony State Office for Consumer Protection and Food Safety and Hannover Medical School, Germany.

#### Isoform gels for myosin isoforms

Different conditions were used to separate and visualize the myosin heavy chains isoforms. To resolve cardiac MyHC isoforms 6. 5 % Acrylamid/Bisacrylamid (100:1) gel was used (66). Stacking gel was prepared using 5 % AA/BisAA containing 5% glycerol. Separating gel contained 6.5 % AA/BisAA with 5% glycerol. The gel was run for 18 hrs at room temperature. The 20 cm long gel (∼4 cm stacking gel and ∼16 cm separating gel) was run in the 1^st^ hour at 10 mA constant and thereafter at 12 mA constant. The gel was stained with quick coomassie stain (Biotrend, Germany). The skeletal muscle myosin heavy chains were separated on 8 % AA/BisAA containing 30 % glycerol. The gel was allowed to run at 30 mA and 4°C for 25 hrs. 12.5 % SDS PAGE was used to separate the light chain isoforms.

#### Preparation of actin filaments

To obtain sufficiently longer biotinylated actin filaments (≥ 20 µm) for optical trapping experiments, chicken G actin and biotinylated G actin was mixed in equimolar ratios to a final concentration of 0.1 µg/ µl each in p-buffer (5 mM Na-phosphate, 50 mM K-acetate, and 2 mM Mg-acetate), containing 1 mM DTT, 1 mM ATP, and 0.5 mM AEBSF protease inhibitor (Cat No. 30827-99-7, PanReac Applichem ITW). The mixture was incubated overnight at 4°C, and followed by addition of equimolar concentration of fluorescent (TMR, tertramethylrhodamine) phalloidin (cat no. P1951, Sigma-Aldrich) and biotin phalloidin (0.23 nM, Invitrogen/Thermofischer Scientific, B7474) to label the actin filaments. For in vitro actin filament motility assays, actin filaments were generated by incubating G-actin in polymerisation buffer (p-buffer) containing 5 mM Na-phosphate, 50 mM K-acetate, and 2 mM Mg-acetate, supplemented with AEBSF protease inhibitor overnight at 4°C. Equimolar concentration of fluorescent phalloidin was added to fluorescently mark the actin filaments. Unlabelled actin filaments were prepared the same way except for addition of phalloidin.

#### In vitro motility assay

In vitro motility assay was performed with full-length βM-II and slow SolM-II by adsorbing the motors on the 1 mg/ml bovine serum albumin (BSA) coated surface. The assay is described in more details in (64). Briefly, BSA containing assay buffer (AB-BSA) (AB; 25 mM imidazole hydrochloride pH 7.2, 25 mM NaCl, 4 mM MgCl_2_, 1 mM EGTA, and 2 mM DTT) was injected into the flow cell and incubated for 5 minutes, followed by myosin infusion. Myosin was allowed to bind on BSA-coated surface for 5 min. Excess/unbound myosin was washed out with extraction buffer. To block the inactive or damaged myosin motor heads 0.25 μM short, unlabelled F-actin was injected in the flow cell and incubated for 1 min. 2 mM ATP was introduced in the chamber to release the actin filaments and to make the active motor heads accessible. ATP was washed out with AB buffer. TMR labelled F-actin was incubated for 1 min, followed by a washing step to remove excess filaments. Finally, the chamber was infused with AB-BSA buffer containing 2 mM MgATP and antibleach system (18 µg/ml catalase, 0.1 mg/ml glucose oxidase, 10 mg/ml D-glucose, and 10 mM DTT) to initiate F-actin motility. For ATP concentration dependence of the motility measurements, different ATP conditions (5, 10, 30, 50, 100, 500, 1000 and 2000 µM) were performed in the same chamber by extensive washing steps in between the two ATP concentrations. Under this experimental set up the surface density of motor molecules were kept constant for motility measurements at low and high ATP concentrations. The fluorescent actin was replenished whenever necessary when multiple ATP conditions were tested in the same chamber. The complete removal of ATP was ensured by observing nonmotile actin filaments in the flow cell, prior to addition of the intended ATP concentration. The sequence of varied ATP concentrations added to the flow cell was changed to ensure the true effect of ATP concentration dependence. This arrangement ensured the identical surface density of myosin molecules and precise effect of ATP concentration on the actin filament gliding speed. Note that the antibleach system was introduced into each ATP containing buffer to ascertain the constant levels of antibleach system as the chamber was used for prolonged period of up to 1 hour, when several conditions were tested. At the end of the measurements, the motility was checked with the same ATP concentration, as at the beginning to confirm that the myosin remained active and yielded similar actin filament gliding speed.

Images were acquired with a time resolution of 200 ms (i.e., 5 frames/sec) using a custom-made objective-type TIRF microscope. Actin filament gliding speed was analysed with Manual Tracking plug-in MTrackJ from ImageJ (NIH, Bethesda, USA).

#### 3-bead assay with optical tweezers

The optical trapping set up was described in detail previously (18,67). For the assay, flow cells with approximately 15 µl chamber volumes were assembled using coverslips with nitrocellulose-coated beads. Glass microspheres (1-1.5 µm) suspended in 0.05 % nitrocellulose in amyl acetate were applied to 18×18 mm coverslips. All the dilutions of biotin-actin filaments were made in reaction buffer (KS buffer) containing 25 mM KCl, 25 mM Hepes (pH 7.4), 4 mM MgCl_2_, and 1 mM DTT. The full-length native myosin was diluted in high salt extraction buffer without MgATP. For the experiment, the chamber was prepared as follows, 1) flow cells were first incubated with 1 µg/ml native myosin for 1 min, 2) washed with high salt extraction buffer without ATP and thereafter with KS buffer, 3) followed by wash with 1 mg/ml BSA and incubated further for 2 min to block the surface, 4) finally, reaction mixture containing 0.8 µm neutravidin coated polystyrene beads (Polyscience, USA) and 1-2 nM biotinylated actin was flowed in with 10 µMATP (or varied concentrations of ATP), ATP regenerating system (10 mM creatine phosphate, and 0.01 unit creating kinase) and deoxygenating system (0.2 mg/ml catalase, 0.8 mg/ml glucose oxidase, 2 mg/ml glucose, and 20 mM DTT). The assembled flow chamber was sealed with silicon and placed on an inverted microscope for imaging and trapping assay.

An actin filament was suspended in between the two laser trapped beads (Figure 2A), pre-stretched, and brought in contact with the 3^rd^ bead immobilized on the chamber surface. Low-compliance links between the trapped beads and the filament were adjusted to about 0.2 pN/nm or higher (26). The bead positions were precisely detected with two 4-quadrant photodetectors (QD), recorded and analyzed. The acto-myosin interaction events were monitored as a reduction in free Brownian noise of the two trapped beads. Data traces were collected at a sampling rate of 10,000 Hz and low-pass filtered at 5000 Hz. All the experiments were carried out at room temperature of approximately 22°C. For the β-cardiac myosin - to improve the time resolution and detect short-lived AM binding events, high-frequency triangular wave of ∼600 Hz and about ± 30 nm amplitude was applied to one of the trapped beads as described in (24,53).

#### Data Analysis

Using the variance-Hidden-Markov-method (27), acto-myosin interaction events detected as reduction in noise were analyzed. This method allowed the AM bound states (low variance) to be distinguished from the unbound states (high variance). Matlab routines were employed to evaluate data records for ‘AM’ interaction lifetime, ‘*t*_*on*_’ and stroke size (*δ*) of motors. Two different methods were employed to measure the stiffness (*k*) of the myosin motors, namely - variance-covariance method and ramp method as described in detail previously (26,27). To calculate motor stiffness with the variance-covariance method, data records with combined trap stiffness in the range 0.05–0.09 pN/nm were used.

Briefly, for the ramp method, a big triangular wave was applied on both the beads. A large amplitude triangular wave of 240 nm and 1 Hz was chosen to study acto-myosin binding events at low ATP concentration (10 µM) due to their long life times, while 120 nm amplitude and 2 Hz wave was applied at high ATP concentrations of 50 µM ATP. Thus, constant ramp velocity of 480 nm/s was used for high and low ATP concentrations. The AM interaction events were registered on both upwards or downwards sides of the ramp. The beads follow the movement of the trap in myosin unbound state, whereas the binding event restricts the bead movement by exerting restraining force, thereby reducing the velocity of bead movements. For the AM binding events, the velocity ratios between bound and unbound states for left and right beads were calculated. The AM cross-bridge stiffness was deduced using the velocity ratios, trap stiffness and combined link stiffness as described previously in Lewalle et al. (26).

#### Ensemble averaging

Ensemble averaging was used to estimate the two-step powerstroke during each individual AM cross-bridge cycle. The computational tool developed by Blackwell et al (25) was used to perform the analysis. To select the AM binding events for this analysis, we define following criterias; 1) the displacement over time record trace from one of the two trapped beads with relatively higher variance ratio (> 5) of bound vs unbound state/free dumbbell noise, 2) the individual events should be well separated, as an indication of single myosin actin interactions, 3) the event lifetime (t_on_) must be more than 50 ms, 4) the binding events (especially the long binding events) trace from both channels/ beads should not contain unspecific interference signal, and 5) the direction of the bead signal fluctuations in both channels is synchronized.

#### Single myosin molecule interaction with actin filaments in optical trapping measurements

For single-molecule optical trapping experiments, we established a protocol to improve the probability that each data record is derived from an intermittent interaction between a single myosin molecule and actin filament. Accordingly, 1) myosin density on the bead surface was adjusted by diluting the myosin solution, 2) among 8-10 beads scanned for the presence of motor on the bead, typically, one bead should interact with the dumbbell, 3) the data traces with distinct well-resolved AM binding events were included in the analysis, 4) closely spaced binding events or stepwise binding indicate multiple molecules simultaneously or consecutively interacting with the dumbbell and such data records were excluded from the analysis. From the Poisson distribution knowing the percentage of beads without motor, we estimate the likelihood of presence of more than 1 motor per bead to be about 5 %. Since we excluded the data traces with closely spaced events, the effective likelihood is less than 5 %. From a total of 432 analysed beads for AM binding events, it is unlikely that a few beads with more than 1 motor could alter our results.

#### Statistical analysis

All values are expressed as mean ± SEM and indicated in the manuscript in relevant sections. Poisson distribution was used to estimate the likelihood of more than 1 molecule interacting with actin filaments in optical trap measurements. Gliding velocities, single motor stroke size, and stiffness of the native and S1 were analyzed using independent samples t-test. The nonparametric Mann-Whitney U test was used to calculate the statistical differences in the duration of AM interaction events, *t*_on_ for the SolM-II and βM-II motors. Statistical significance was assumed if P < 0.05.

## General

We are grateful to Petra Uta for the technical assistance in protein preparations, Faramarz Matinmehr for help with the selection of type-1 muscle fibers, and Ante Radocaj for critical comments on the manuscript. We thank Walter Steffen (deceased) for his crucial inputs in the experiments.

## Author contributions

MA conceived the project and designed the experiments with inputs from TW, and AN. TW performed optical trapping measurements and analyzed the data. MA performed motility experiments, and ES, JO, FB, FG, TS, performed data analysis. Motility and single molecule data was interpreted by TW, AN, and MA. TS and TK were involved in the discussions. MA wrote the manuscript with assistance from TW and AN. All the authors contributed to the editing of the manuscript.

