## Supplemental Information for "Cardiac ventricular myosin and slow skeletal myosin exhibit dissimilar chemo-mechanical properties despite the same myosin heavy chain isoform"

**This file includes:**

(1) Figures S1 to S3:

Figure. S1. Myosin isoforms gel for left ventricular myosin.

Figure. S2. Myosin isoforms gel for soleus myosin.

Figure S3. Stiffness of βM-II.

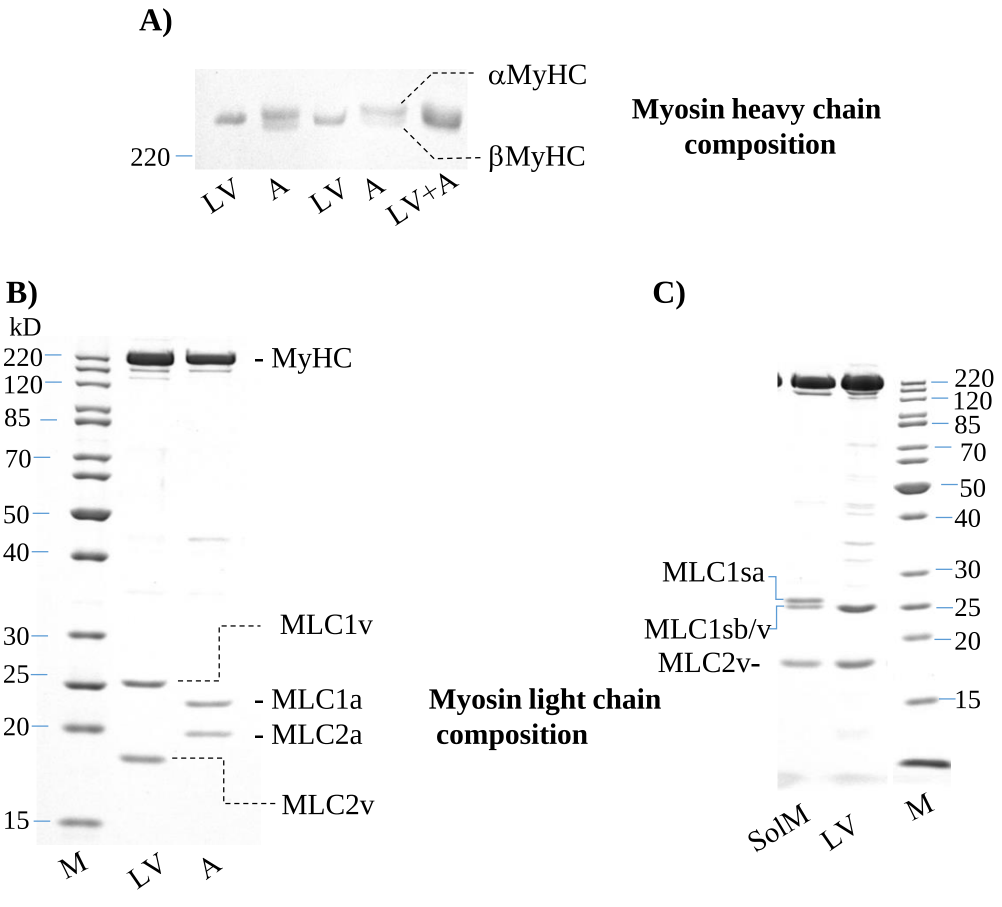


**Figure S1. Myosin isoform gel for βM-II**. Heavy and light chain composition among α- and β- myosin II as well as M. Sol myosin and β- myosin compared. **A)** Myosin II isolated from rabbit left ventricle (LV) and atrium (A) tissue. 5 % acrylamid/bisacrylamide gel with 5 % glycerol was used to separate the α- and β- myosin heavy chain isoforms. The long gel (20 cm) was run for 18 hrs at room temperature and followed by coomassie staining. For lane 1 and 2 - 0.5 µg of protein sample, lane 3 and 4 – 0.375 µg sample, and for lane 5 – mixture of 0.5 µg each of atrial and ventricular myosin was loaded. While, two myosin heavy chain isoforms (α-MyHC and β-MyHC) as two distinct bands were detectible in atrium derived myosin, ventricular tissue showed only single band corresponding to β-MyHC. Note that below 5% contamination of α-MyHC in the ventricular tissue derived myosin is unlikely to be evident in the gel analysis. B) The light chain compositions were analyzed on 12.5 % gel. Distinct banding pattern corresponding to the essential and regulatory light chains typically associating with the α-MyHC and β-MyHC were observed. αM-II and βM-II, in each lane - 2 µg protein is loaded to check the light chain composition. C) Light chain composition of LV myosin and M. soleus myosin (SolM). 4 µg of proteins loaded in each lane.

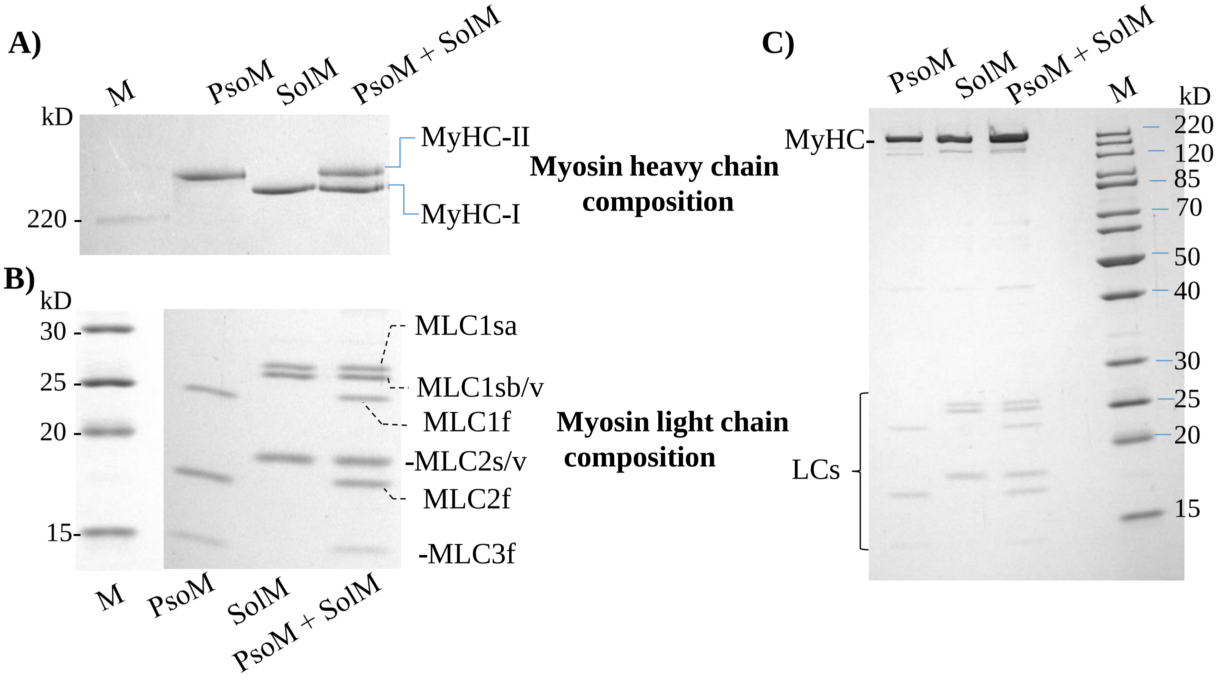


**Figure S2. Myosin isoform gel for SolM-II. A)** 8% acrylamid/bisacrylamide gel with 30 % glycerol was used to separate the myosin heavy chain isoform. The 20 cm custom-made gel was run for 25 hrs at 4°C. 0.5 µg of protein/lane was loaded. Lane 2 and 3 – isolated protein from two muscle sources M. psoas (PsoM) and M. soleus (SolM) . Lane 4- mixture of protein (0.5 µg each) used in lane 2 and 3 were loaded. Fast M. psoas myosin heavy chain (MyHC-II) is seen higher than slow M. soleus myosin (MyHC-I) with distinct banding. **B)** Same probes as in A) examined for corresponding light chains for the two myosin isoforms on 12.5 % SDS-PAGE gel. Distinct light chains, i.e., MLC1sa (27 kD), MLC1sb/v (24 kD), and MLC2s/v were seen to assemble with the SolMII, whereas PsoMII can be found in complex with MLC1f (20.7 kD), MLC3f (16.5 kD), and MLC2f. **C)** 12.5 % gel for the same sequence of myosins loaded showing both heavy and light chains. Note that the heavy chain isoforms cannot be distinguised in this gel. LCs- light chains, M- marker.

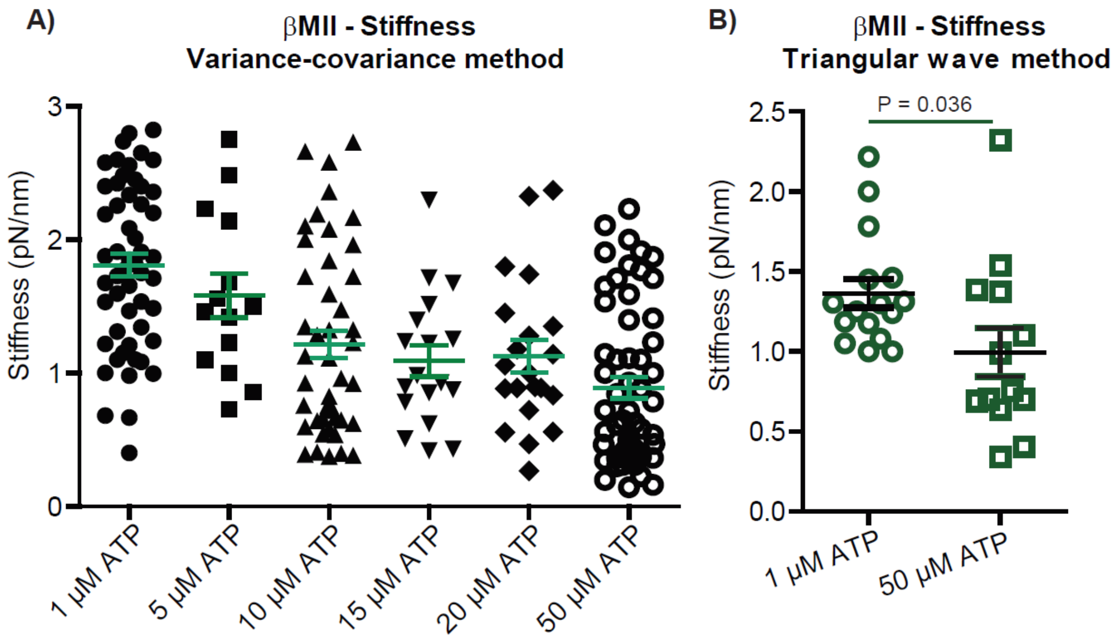


**Figure S3. Stiffness of βM-II. A)** Stiffness of individual βMII molecules measured at various ATP concentrations shown. Each data point indicates the average stiffness determined from 100s of AM binding events for individual myosin molecule. Myosin Stiffness estimated using Variance-covariance method. **B)** Myosin stiffness measured at low (1 µM) and high (50 µM) ATP concentration. 1 µM ATP; N= 17, n= 802, 50 µM ATP; N= 13, n = 301, N = number of molecules, n = number of events.
